# Defining the molecular tolerance-to-damage landscape of SMARCA4 helicase genetic alterations

**DOI:** 10.64898/2026.06.22.733755

**Authors:** Neshatul Haque, Xiaowei Dong, Jessica Wagenknecht, Michael T. Zimmermann

## Abstract

Evaluating the impact of genomic variation is essential for identifying underlying mechanistic causes of human diseases. The spectrum of neurodevelopmental disorders is driven by diverse genetic alterations with genes like SMARCA4 being prototypical examples. There have been significant hurdles to implementing the protein-specific and mechanism-informed variation effect predictors that are anticipated to have the highest yield of mechanistic information. Yet, there is a pressing need, for example, within SMARCA4 where 98% of the 2780 reported variants lack a disposition and remain of uncertain significance (VUS). Further, the field has yet to identify each variant’s specific molecular mechanism, which will inform targeted therapeutic development strategies. In this study we developed a mechanistic structure-informed helicase-specific variant effect predictor by leveraging diverse information with state-specific calculations. Our approach has 100% recall of pathogenic variants while classifying 87.23% of VUS into damaging (55.74%, n=262) versus tolerated effects (31.49%, n=148), including those with conflicting interpretations. This analysis reveals significant enrichment of integrated functional metrics, such as conservation and solvent exposure, that parallel allele frequences in health populations, and emphasizes the robustness of the method. Thus, we have demonstrated a novel approach for the development of mechanism-informed protein-specific interpretation of human genetic information.

## Introduction

The BAF complex is well studied master architect of the cellular landscape. It regulates precise spatiotemporal accessibility of the genetic code by sensing environmental signals defined by epigenetic signatures (3, 4). BAF complexes are comprised by combinations of 30 different genes in lineage-specific manners, with the canonical configuration of 12 genes (1, 2, 5). SMARCA4 is one of the two possible core ATPase engines of all BAF complexes. SMARCA4 generates the chemical-to-mechanical force for moving the DNA wrapped around histones. When the coordinated motion of the BAF complex is disrupted by genetic variations in SMARCA4, the functional change can be severe, resulting in rare diseases or malignancies. For instance, genetic variations in the SMARCA4 helicase are linked to developmental disorders like Coffin-Siris (CS) syndrome (6, 7) and aggressive cancers such as Small Cell Carcinoma of the Ovary, Hypercalcemic Type (SCCOHT)(8). While SMARCA4 is a tumor suppressor, specific DNA-binding amino acids are more often the site of substitutions (9) resulting in modest somatic hotspots. However, there are many more people affected by the diverse inherited and *de novo* variations underlying congenital neurodevelopmental syndromes (6, 7), plus somatic alterations across the gene body (8–10), compared to these modest hotspots. Currently, 2780 distinct missense genetic variants have already been observed in human specimen, representing 8.88% of the theoretical maximum of all possible protein alteration in this region. Interestingly, 2726 (98.00%) of observed missense genetic variations are classified as VUS. and are typically ultra-rare (96.36%). Therefore, new methods are needed to scale mechanistic interpretation of diverse and ultra-rare variation.

Before variant effect modeling, variant effect prediction (VEP) has been a challenge in modern medicine. Myriad VEPs have been trained on massive protein sequence data (11–17), augmented with structural features predicted from sequences (18–23), and integrated with genomic features or combined into meta-predictors (15, 24–30). While high-throughput VEPs effectively estimate the global pathogenic likelihood of genomic variants, their generalized architecture lacks the resolution required to interpret mutations at protein-specific functional residues. Hence, these models fail to account for the unique physicochemical constraints or specialized catalytic roles, interfaces compatibilities, or binding site affinities characteristic of individual protein domains and required for their normal cellular and physiological functions. Such limitations are inherent to sequence-based approaches which led to the development of structure-based variant effect predictors (31–37). The progression of protein sequence-based models presents a compelling opportunity and solid groundwork for developing protein-specific models that account for the precise and accurate molecular features of individual proteins. The nuanced and critical difference is that long-standing science has demonstrated that proteins are robust to mutation (38, 39), yet specific point substitutions cause human diseases. Distinguishing lynchpins from the rest remains a challenge. In this way, further development of more structurally-informed VEPs will improve variant effect modeling.

In this study, we used structural properties to investigate the most likely origin of damaging effects for all observed human genetic variation across the SMARCA4 helicase domain. The helicase domain is comprised of two sub-domains, SnF2 and HelicaseC, according to sequence conservation and evolution. Its mutational landscape is consistent with its tumor suppressor role - a consistently low-level variation across the full length of its linear sequence, with limited regional differentiation and a few modest peaks (i.e., monadnock-strewn). These isolated peaks often cluster in close spatial proximity. The 3D clustering of these functional residues underscores their biological significance and likely coherent functional mechanism. Our new data suggests that helicase domain variants most frequently impact the scaffold stability (51.30%), substrate binding (4.50%), state transitions (3.60%), and ATP binding (0.76 %). Further, there is a hierarchy among these molecular mechanisms such that genetic variants that compromise overall fold stability are more likely to also impair DNA binding or be incorporated into chromatin remodeling complexes. Thus, foundational instability obviates the other stages of helicase function. Therefore, we present evidence that a model tailored to the SMARCA4 helicase domain will yield a more robust and reliable variant effect prediction compared to models trained on the proteome.

We propose that domain-specific models augment genome-wide or proteome-wide predictions such that general-purpose VEPs setup the context within which our protein-specific calculations should be interpreted. In our approach, the power of genome-wide VEPs is combined with protein specific features calculated using specific context-aware structures. Further, we leverage multistate structural features complete with mechanism-informed functional assignment on the residue level. Our work yields a substantial gain of rationally defined molecular mechanisms ascribed to 87.24% of variants, representing a potential 7.60-fold increase of interpretation yield compared to the current state. We anticipate that data of this type is highly useful for addressing VUS ambiguities for individual patients, and future therapeutic investigations towards the applicability of inhibitory and degrader technologies across constitutional and somatic diseases. Therefore, the mechanistic investigation of human variation will undoubtedly play a critical role for illuminating the underlying causes of genetic diseases and enable a new phase of human genetics research, towards the interpretation of individual genomes complete with mechanistic variant effect modeling.

## Methods

### Genetic Variation Information and Annotation

SMARCA4 helicase missense variants (Figure 1A) are obtained from ClinVar (40, 41), GnomAD v4 (42), and UniProt (43) variant databases. Pathogenicity prediction scores are obtained from dbNSFP(44) v5.0a . Germline variants were annotated using pathogenicity prediction tools including REVEL(30), CADD(24, 25), SIFT4G, PROVEAN(12), ESM1b(13), AlphaMissense(18), DANN(26), Polyphen2(19), Eigen(27), MutationAssessor(45), MetaLR(28), fathmm(14), DEOGEN2(20), HACTboost(21), ClinPred(29), MutFormer(17), MutScore(23), VARITY(22), PrimateAI(16), and BayesDel(15). We calculated the cross-correlation among 39 pathogenicity scores. Scores with mutual Spearman correlation above the threshold of 0.85 were filtered, leaving 20 sequence-based features for subsequent analysis.

**Figure 1:**
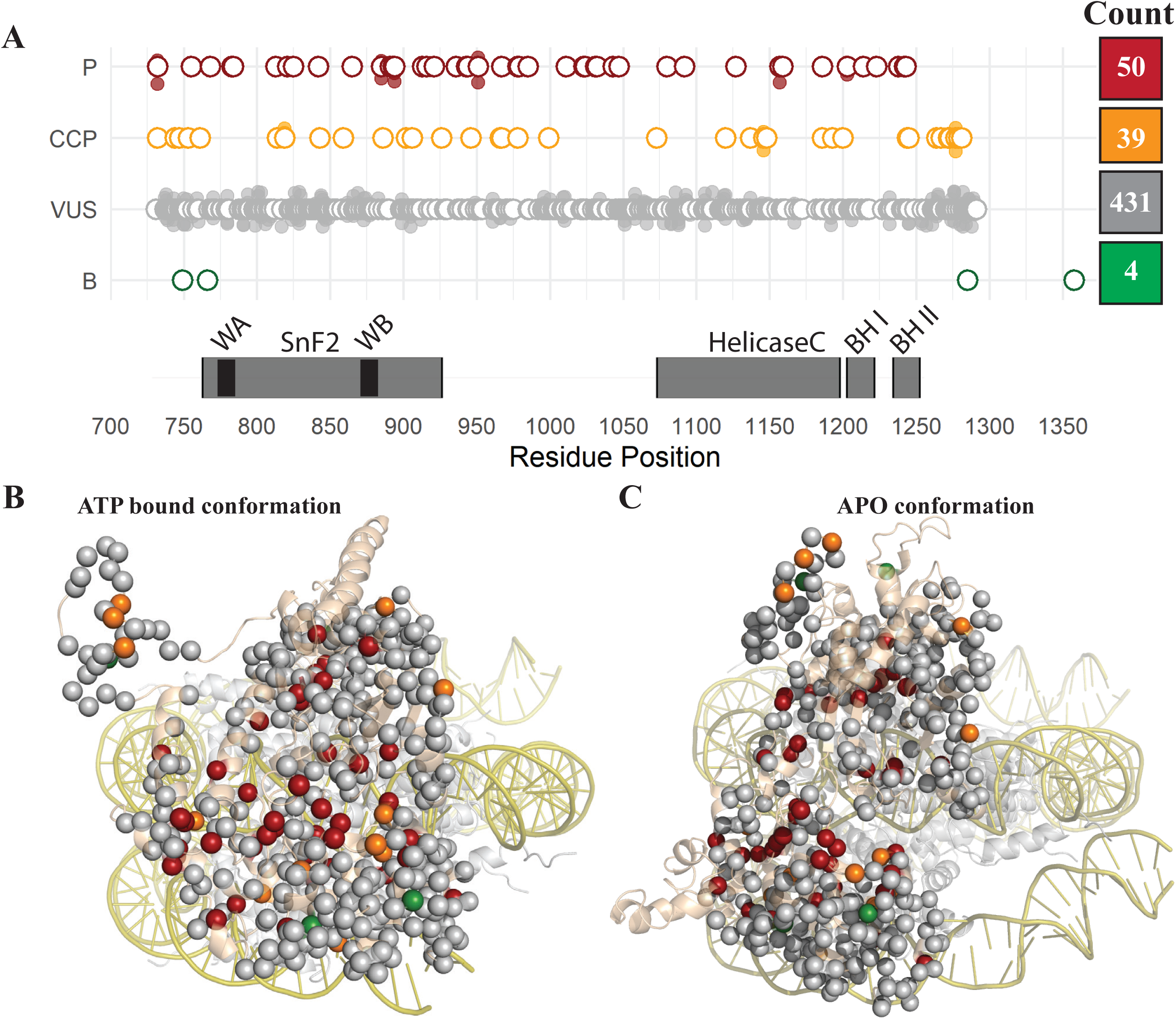
SMARCA4 genetic variations frequently affect the bipartite helicase. **A)** ClinVar variations occur across the entire helicase sequence, among all classification categories. Open circles mark the first variant at each location. When two or more variants occur at the same location, they are marked by filled circles and offset. Counts of each variant category are shown on the right. The bipartite helicase domains, ATP-binding (aa766-931) and HelicaseC (aa1084-1246), are marked as domain blocks. Additionally, some of the functionally critical regions such as, Walker A (WA), Walker B (WB), Brace helix I (BH I), and Brace helix II (BH II) are also marked. Helicase-Nucleosome 3D structures were obtained in **B)** ATP-bound from (PDB: 7VDV (1)) and **C)** Apo (9A0K (2)). A semi-transparent rectangle shows clear separation between the two domains, at the nucleotide binding region.

### Protein Structural Characterization

The structures of SMARCA4 helicase in ATP bound(2) (Figure 1B) state and Apo states(1) (Figure 1C) were selected for calculating differences in stability metrices, 3D clusters, and chemical interactions at the residue level. The structural stability of the ATP-bound and Apo state were estimated using FoldX(46) v5.0 and for generating mutated structures. We used frustratometer(47) v0.1.0 for calculating structural energy of variants, and STRIDE(48, 49) for residue surface areas. The relative solvent accessible surface area is calculated using maximum surface area data from naccess(50) v2.1.1. Collectively, these tools provided 14 structural stability scores, nine from ATP-bound and five from the difference from ATP to Apo conformational states, encompassing energetic parameters such as total and solvation, configurational, mutational energy relative to the wild type, and representing the basic thermodynamic cycle.

### Developing an Integrated Variant Assessment Procedure for SMARCA4

Currently, clinical interpretation largely relies on co-observation of genetic variation in affected versus unaffected individuals. Since SMARCA4 is missense intolerant in the germline (z = 8.81) (42), there are relatively few variants available as confident negative controls. Additionally, while there are many clinically observed VUS, there are only 4 that have reached the level of (likely)benign and 50 (likely)pathogenic and are therefore available as negative and positive markers. Due to the scarcity of clinically annotated data, we sought a data-driven means of comparing variants in an all-versus-all framework. We developed an unsupervised learning approach utilizing K-means and Gaussian Mixture Models (GMMs) applied to combinations of 20 genomics and 14 structural features. First, each of the 34 features were normalized into z-scores. Clustering was then performed on both the normalized data and on the first 10 principal components (PCs) derived from the normalized data. The first 10 PCs collectively accounted for 81.3% of the total variance. The combination of two input types (untransformed and PC) with two cluster methods (KM and GMM), resulted in four approaches to group variants by their relative similarities. We identified that one cluster captured all (likely)pathogenic variations, while the second cluster contained 75% of benign. Thus, we chose to use clusters to classify damaging from tolerated alterations to the SMARCA4 helicase (Figure 2).

**Figure 2.**
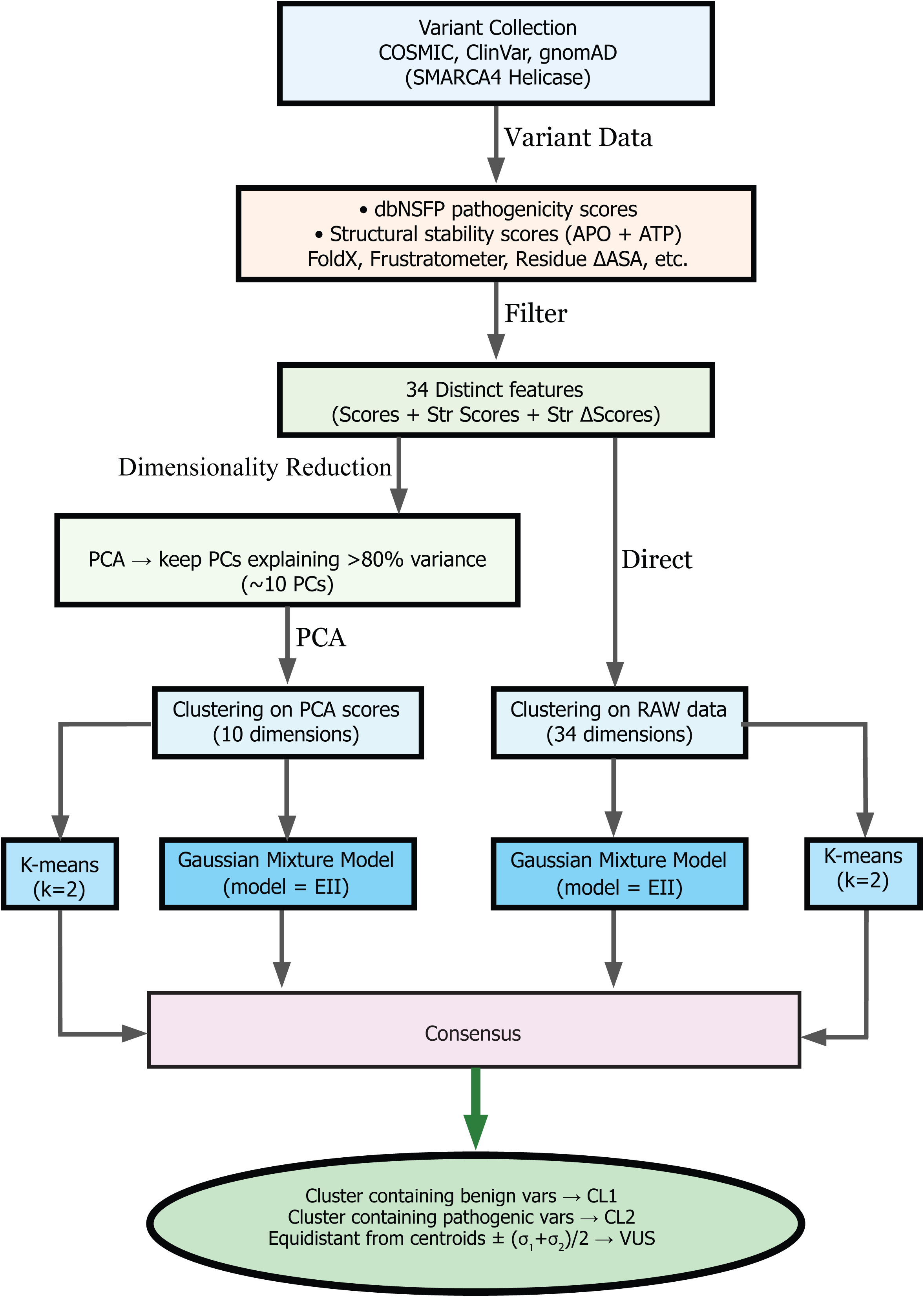
SMARCA4 helicase variant reclassification procedure based on structure-based calculations. Each major step implemented in this study is shown in their order of operations, with key parameters used.

For each clustering approach, we calculated each data point’s distance to cluster centers (C_1_ and C_2_) (using respective dimensions of the data as column mean, **µ_1_** and **µ_2_**and standard deviation, **σ_1_** and **σ_2_**) and obtained the difference of the distances (d_1_ and d_2_), denoted by Δd = d_1_ – d_2_. The clustered variants were then strictly evaluated for cluster membership (Figure 3). This approach gives us power to accurately classify damaging from tolerated variation by similarity to (likely)pathogenic and (likely)benign, with variants that remain ambiguous or variable in clustering remaining as VUS. We assessed cluster ambiguity using an exclusion method where variants near the midpoint of the two cluster centers, defined in units of pooled standard deviation according to |Δd| ≤ |(**σ_1_** + **σ_2)_/2**| and for each approach, were marked as VUS (Figure 3). The consensus model is developed with the rules described in Table 1 and used to reclassify all the ClinVar VUS and variants with conflicting classification of pathogenicity (CCP). Results obtained were subjected to statistical tests such as Pearson’s chi-square, Likelihood Ratio (G), and Fisher’s Exact test for testing the enrichment or depletion on the distribution of functional metrics, such as rSASA and the evolutionary conservation of mutation sites, presence of variants at interfaces, and variant ascertainment source like gnomAD(42), and TCGA(51).

**Figure 3.**
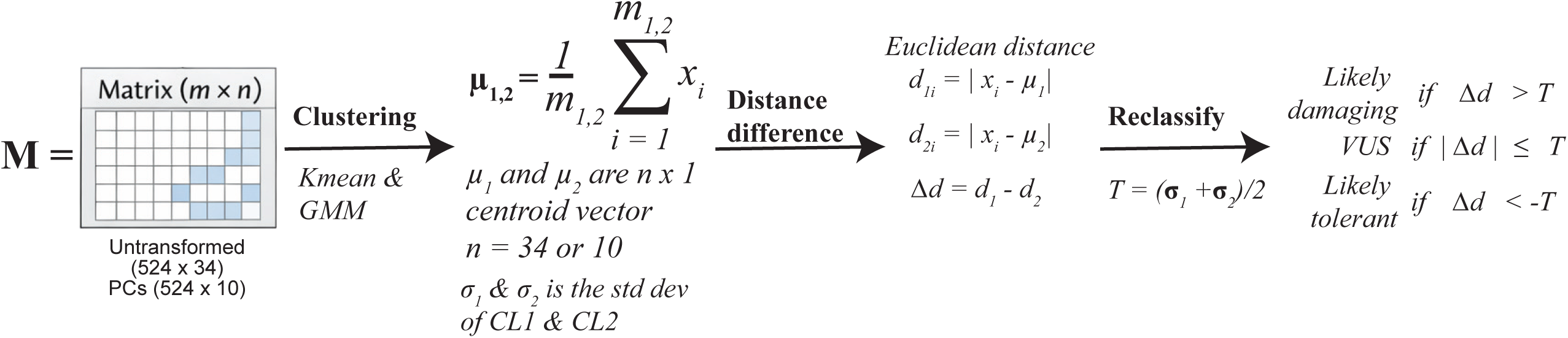
Exclusion based reclassification method: The flow diagram represents the overall process of clustering (by K-means and GMM) followed by reclassification.

**Table 1.** Rules implemented on the four models to develop the consensus model.

| Situation | Consensus | Rationale |
| --- | --- | --- |
| Variant is established as B | B | Gold standard from clinical observations |
| Variant is established as P | P | Gold standard from clinical observations |
| $\geq 1$ method, "Ambiguous" | VUS | Conservatively retain uncertainty |
| $\geq 3$ methods, "Likely tolerant" | Likely tolerant | Clear consensus majority |
| $\geq 3$ methods, "Likely damaging" | Likely damaging | Clear consensus majority |
| Exactly 2 tolerant + 2 damaging | VUS | Disagreement maintains uncertainty |

## Results

### Landscape of SMARCA4 helicase missense genetic variation

SMARCA4 is a core component of the epigenetic regulatory system. As such, it has profound constraints against developing genetic variations, evidenced by high missense intolerance (z = 8.81) and maximal pLI (probability of Loss-of-function Intolerance) = 1.00. Interestingly, SMARCA4 genetic variations are observed frequently in neurodevelopmental rare diseases and cancers from across many body systems (10, 52). Loss-of-function alleles are regarded as easier to interpret, with assumed uniform loss-of-function, while missense presents a more nuanced challenge. Therefore, we obtained all observed (n=524) missense germline variations within the SMARCA4 helicase from across constitutional and somatic diseases and healthy populations (40, 42, 43). We observed that most of the cancer associated variants are rare or ultra rare (2.8%, and 23% of n=524, respectively) and very few are common in the healthy population (1.14% of n=524). Disease-associated and non-disease-associated variants often occur within key functional sites, such as Walker motifs and Brace helices (Figure 1A), or line the interfaces of the domain interaction. Thus, structural calculations are likely to help distinguish among tolerated and damaging variants.

The primary function of the SMARCA4 helicase stems from its capacity to transduce chemical energy from ATP into mechanical energy that separates and moves DNA relative to histone proteins. To accomplish that task, multiple components must align, chemically and spatially. In this way, the fold architecture itself is needed to scaffold chemical groups from catalytic amino acids with interface surfaces. The stability of the fold, and therefore of the scaffolding functions, are of critical importance. The scaffold stability of the SnF2 and HelicaseC domains, enzymatically critical motifs like Walker A and Walker B, and transition state stabilization characterized by regulatory helices including Brace I and Brace II. During helicase catalytic cycling, Brace helices appear to undergo order-disorder transition upon nucleosome binding (1, 2, 53, 54) to physically brace the two lobes of the helicase enzyme. When Braces are disordered, the two lobes of the helicase are more separated in space, concordant with Apo conformation and differences in DNA binding compared to ATP-bound. We anticipate that alteration of these properties, such as by genetic mutations, can affect the overall cycle of nucleosome binding, ATP hydrolysis, and post hydrolysis transitions, leading to change in helicase function and subsequently the BAF regulated chromatin remodeling. Studying these variations will help understand their molecular role, generating opportunities for mutation specific therapeutic design and treatment strategies.

### Current clinical classifications of SMARCA4 helicase missense genetic variation

SMARCA4 helicase is the ATPase enzyme responsible for generating and transducing chemical-to-mechanical force transduction required for torque generation during nucleosome remodeling. Most of the variants are categorized as VUS (n = 431, 82.25%), followed by (likely)pathogenic (labeled as P for simplicity, n = 50, 9.54%), conflicting class of pathogenicity (CCP, n = 39, 7.44%) and (likely)benign (B, n = 4, 0.76%). Most (n = 32, 64%) pathogenic variants occur within the ATP binding (aka, SnF2-like) and C-terminal helicase domains (Figure 2A-C). We calculated the proximity of variants to key functional sites in 3D (Figure 1B-C) and found that 52% of pathogenic variants (n = 26) are proximal to the ATP binding site. Because the domains are well-separated in the Apo state, we can clearly differentiate between pathogenic variant clusters at the interface and active site, which are adjacent and therefore appear as one overlapping group in the closed ATP-bound conformation. Therefore, it is evident that functionally critical, and thus mutation sensitive, regions of helicase are frequently populated by pathogenic variants, suggesting that the dysfunction of two domains is a central driver for nucleosome remodeling diseases.

### Classifying genetic VUS by co-similarity across multi-state structural features

Seeking to scale the tolerance-to-damaging landscape, we leveraged high-dimensional similarity measures to split the variant dataset into three categories based on clustering. First, we calculated relative similarities among the full collection of missense variants. Then, we clustered the variants into two groups, testing for cluster consistency and separation. Next, we identified that data-driven clustering already achieved 100% recall of the known (likely)pathogenic variants and a WT-adjacent region that co-cluster 3/4 benign variants. It is often a challenge to identify benign controls because of the rigorous evidence required to classify them. Specifically, the two most common criteria set are to definitively prove a lack of functional impact in patient-derived materials using a defined and accepted functional assay or demonstrate sufficiently high prevalence in the general population in contrast to affected patients. In this way, we then projected VUS onto an in-common metric space and re-classified them according to their high-dimensional similarities. We propose evidence supporting that most of the VUS (n = 343, 65.4%) be re-classified as LD, while about a third (n = 169, 32.2%) be re-classified as likely tolerated.

Clustering results from the four approaches are highly concordant (Figure 4A), with pairwise classification agreements approaching 98% (Figure 4B). This suggests that the methods used are effective in categorizing missense variants. Thus, a structure-based data-driven approach can potentially enhance protein specific variant classification to better interpret inter-individual genetic variation.

**Figure 4:**
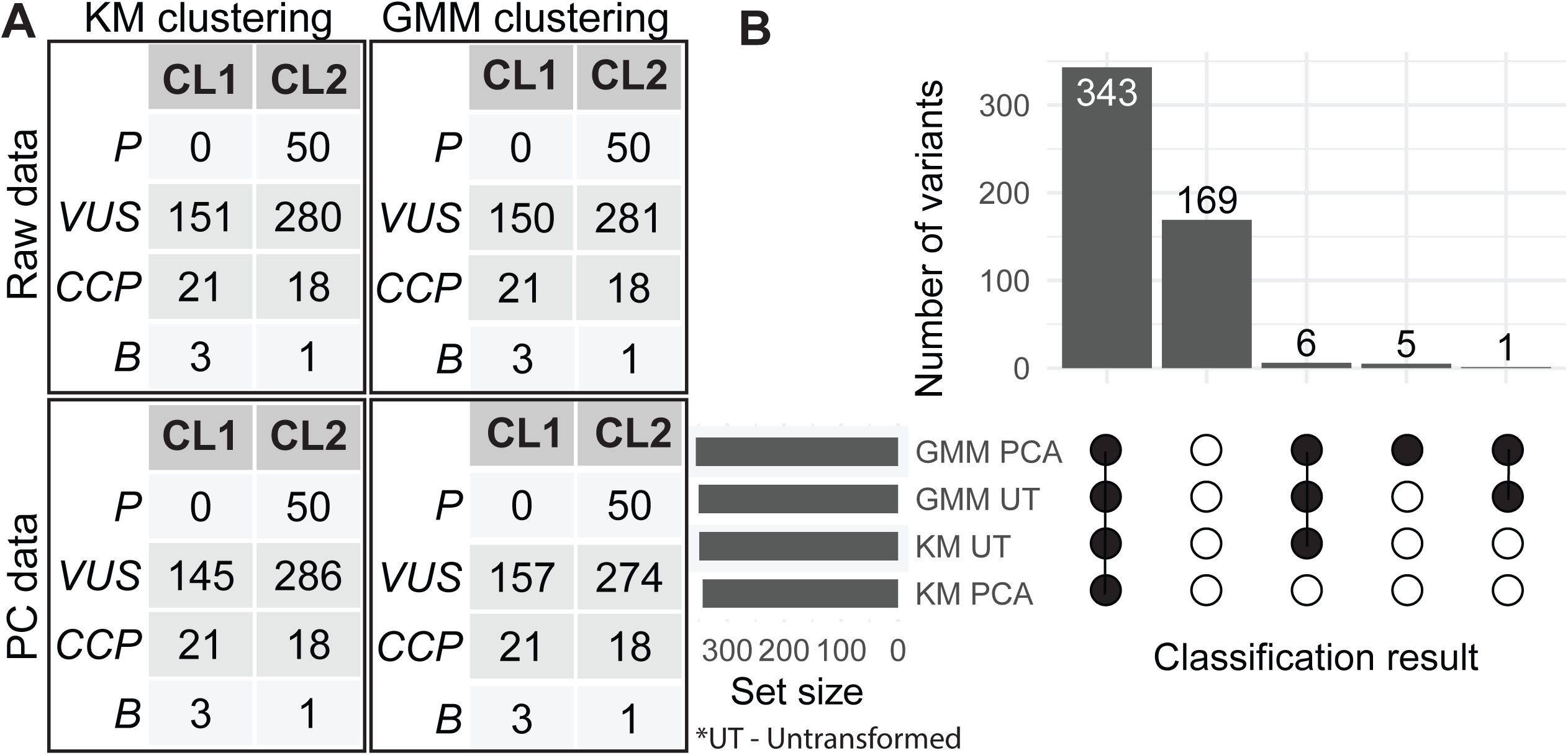
Variant re-classification is robust to implementation parameters. **A)** Contingency tables across normalized (34 features) and PC-based (10 features) datasets on 524 variants, using either K-means or GMM clustering. **B)** Upset plot of the pathogenic-associated cluster membership shown in black circles and benign-associated cluster shown in white circles.

**Figure 5:**
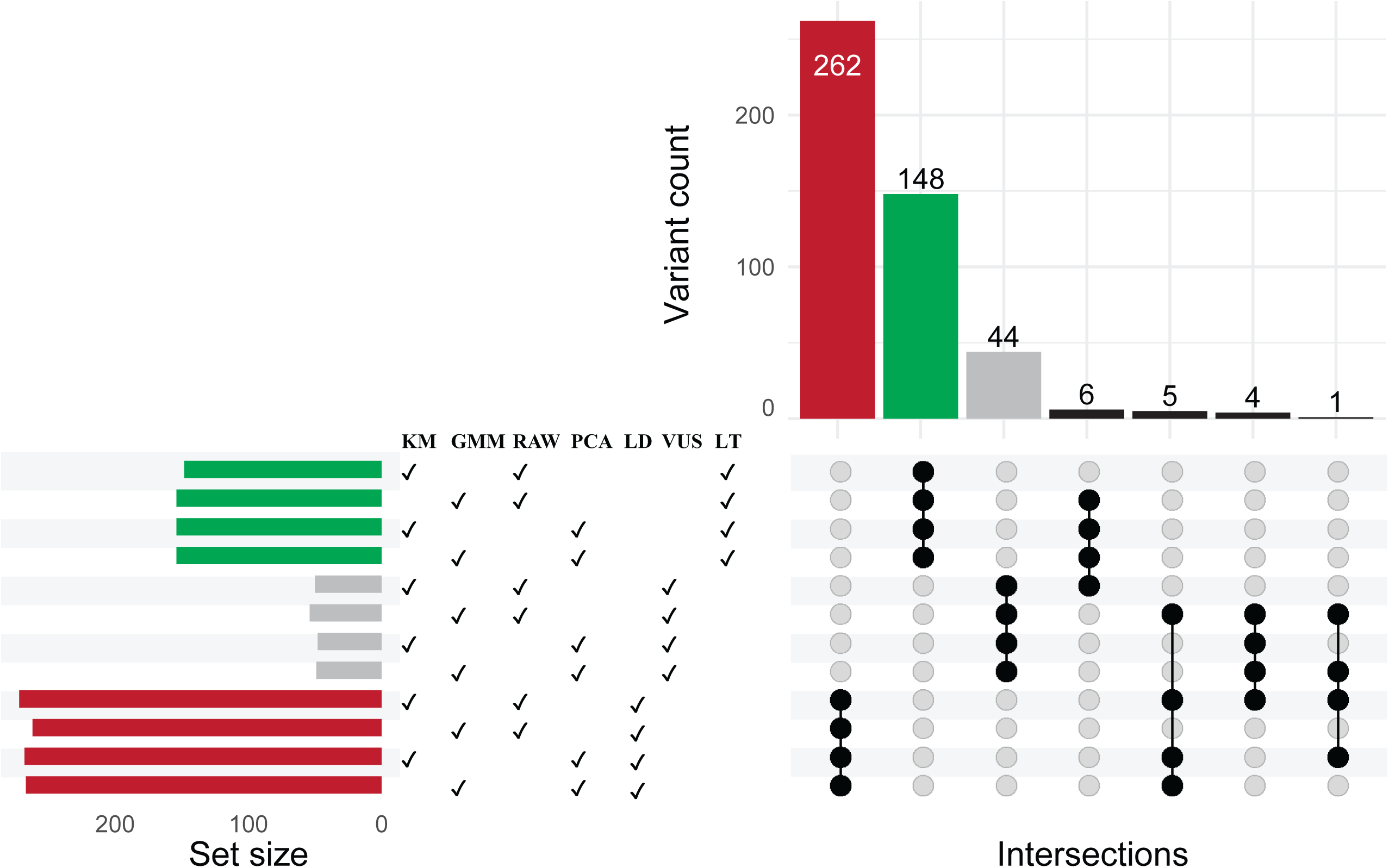
Reclassification is highly consistent and retains ambiguous variants as VUS. The distribution of variants upon reclassification is shown as an upset plot where check marks indicate which input data types and clustering algorithm were used. Cluster 1 contains ClinVar benign and, cluster 2, ClinVar pathogenic variation.

Following our re-classification, we applied a second approach to further increase robustness. This exclusion-based cluster membership assessment (Figure 3) used the distance difference (Δ*d* between variant feature vectors and each cluster’s central point (Figure S2A). The equidistant region (Δd ≈ 0) effectively demarcates the boundary between the two clusters. The subsequent projection of Δd on the three-dimensional protein structure model highlights a high incidence of likely damaging variants at the interface between domains and within each domain’s folded core. Concordantly, likely tolerated variants were observed more often along the spatial periphery. The equidistant region (Δd ≈ 0) was observed to have a mixed distribution of spatial features (Figure S2B), consistent with the group of variants having moderate or divergent effects on the same helicase features as pathogenic variation. The biological relevance of these distributions are more evident when mapped jointly onto ATP-bound and Apo states. The mechanism driving variant class discrimination is a conspicuous nucleotide-dependent separation between the domain lobes. Leveraging information between both states increased the consistency of re-classification results. Re-classification yields 50% of current VUS predicted as likely damaging (LD), 28% remaining as VUS, and 8.4% predicted as likely benign (LB) with only 3.2% discordant. Consequently, the observed consistency in clustering confirms the discriminatory power of enhancing genomics scores with structure-based calculations. The features we used effectively classify variants based on quantitative physico-chemical properties of the residues that emphasize their functional roles.

### Consensus classification yields robust outcomes

To further quantify how our consensus classification coincides with physical features that are established as correlating with structural damaging effects, plus if our methodology is robust to differences in clustering and distance measures, we implemented a rubric for consensus classification leveraging all previous clustering results (**Table 1**). We used dimensionality reduction to enable visualizing the high-dimensional relationships involved (Figure 6A, Figure S1). Consensus results identify unannotated variants that are likely damaging to the helicase fold (56.59%) and likely tolerated by the fold structure (32.13%). The remaining 11.27% of variants fell within the consensus ambiguous region, potentially indicating that they may have mild, atypical, or divergent effects on the fold structure. We chose to retain them as VUS (Figure 6B). In this way, we propose a 78.24% increase in the number of dispositioned variants by leveraging structural bioinformatics for mechanistic interpretation.

**Figure 6:**
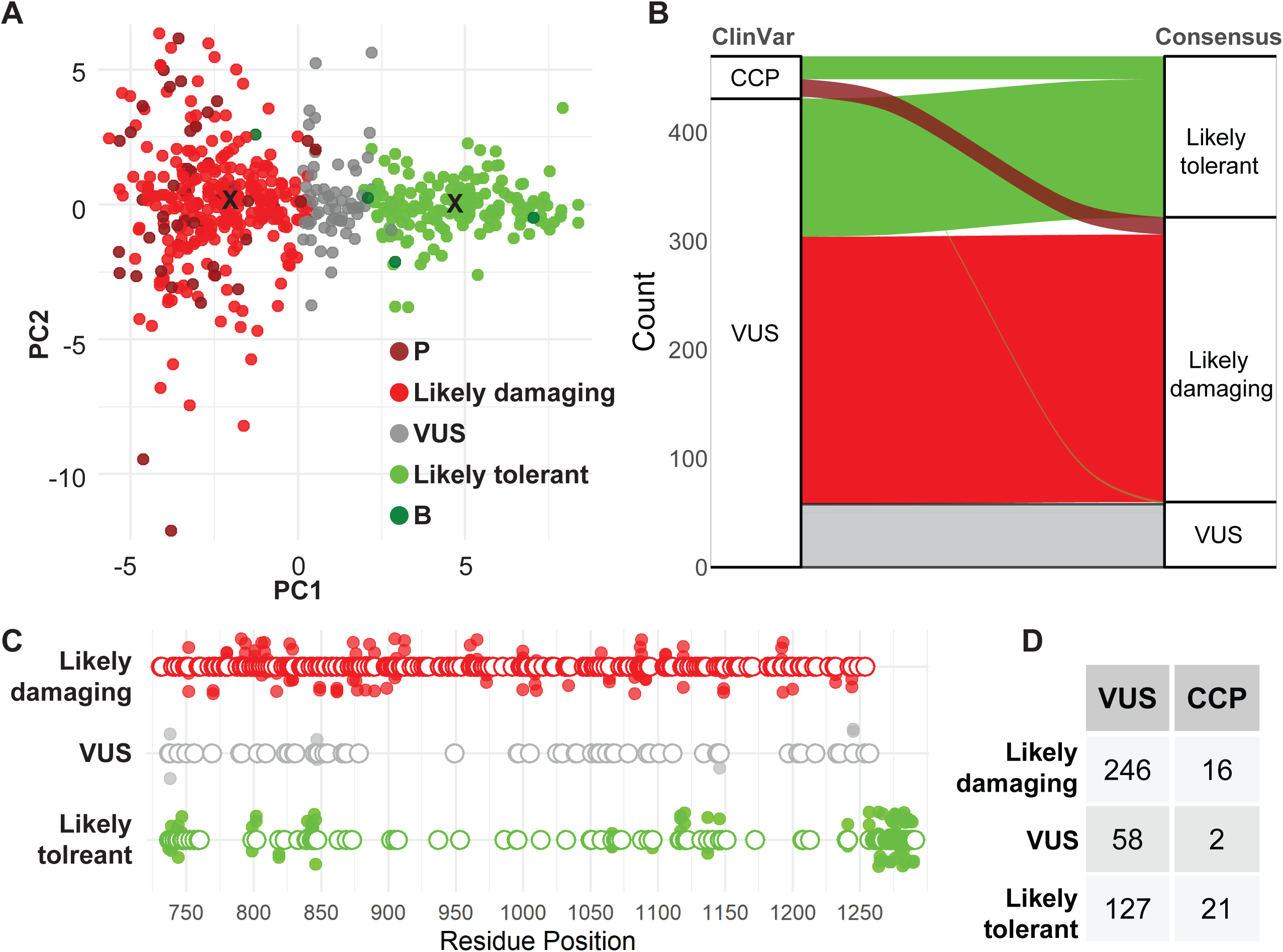
Consensus classification generates ensemble model: **A)** For visualization of the structurally reclassified variants, PC1-PC2 scatter plot shows where the P and B dominated clusters occur and their relationships to variants equidistant from both the cluster centroids (marked with “x”) that remain VUS. **B)** Visualization (Sankey plot) of redistribution of CCP and VUS in Three categories of Consensus class **C)** Linear mapping of reclassified variants on helicase. **D)** Contingency table of Consensus classification for ClinVar VUS and CCP.

### Integrating functional and physicochemical information for class interpretation

We assessed our consensus findings by calculating enrichment and depletion metrics reflecting physical and chemical properties of each mutated residue’s functional state. We evaluated several functional metrics including how much relative solvent accessible surface area differs for the mutation and between nucleotide-dependent conformations of the helicase (e.g., rSASA), and the evolutionary conservation of mutation sites. Additionally, we characterized the distribution of variants at the chemically active motifs, domain-domain, and domain-DNA interfaces. Then, we also performed a comparative analysis of variant prevalence in healthy population and oncology-specific database (The Cancer Genome Atlas, TCGA). The resulting data suggest that the (likely)pathogenic variants are distributed mainly in the interior of the helicase domain (rSASA < 33%). We found that buried residues are enriched in LD and depleted in LT class. Exposed residues are depleted in LD and enriched in LT (p-value =2.00×10^-16^). Residues at the interfaces are also crucial for helicase function as they participate in regulating the strength, efficiency, and timing of the BAF complex’s mechanical dynamics. Our analysis further reveals a significant enrichment of known pathogenic variants at the interface, whereas such variations are notably depleted in other regions. Similar observation was noted for LD and LT (p = 1.9×10^-4^). Population variation coincides with and is statistically enriched in the LT category and depleted among LD (p = 3.5×10^-4^). Cancer-associated variation is enriched in LD and depleted in LT, (p = 2.3×10^-5^). Conserved residues were enriched in LD and depleted in LT (p = 6.2×10^-3^) (Table S1 Figure 7 and S3). Thus, the damaging-to-tolerated landscape of SMARCA4 helicases follows expected patterns for functional protein variation, supporting our application of these advanced scoring rules to interpret human genetic variation.

**Figure 7:**
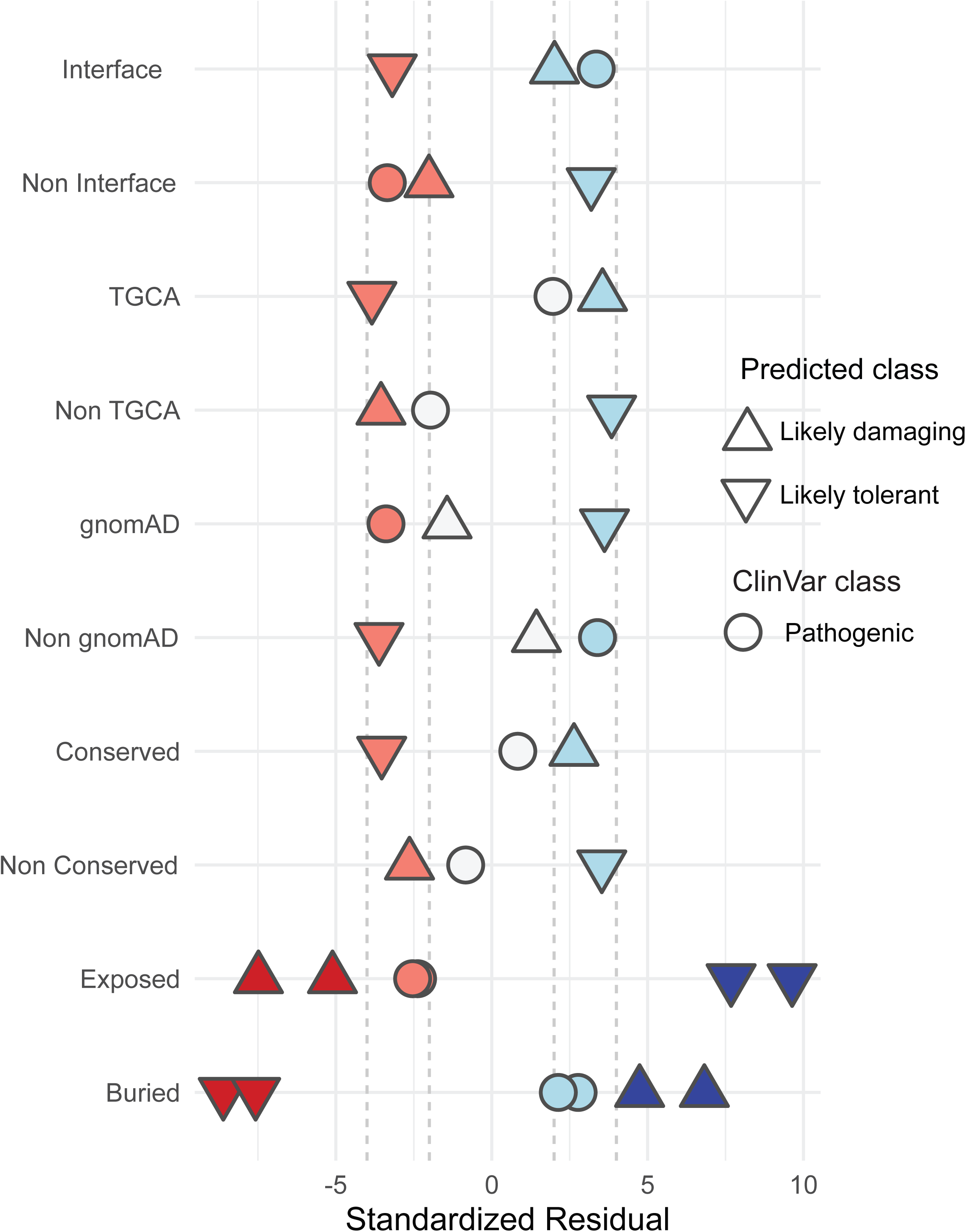
**Statistical enrichment of functional metrics across variant categories**. Plot values represent Adjusted Standardized Residuals (Haberman’s residuals), which follow a standard normal distribution (z-scores). Residuals falling outside the −2, 2 range indicate rejection of the null hypothesis and are statistically significant (p < 0.05). Values exceeding +2 represent significant enrichment while below −2 represent significant depletion.

### Pathogenic variations change physicochemical properties consistent with loss of function

Pathogenic variants are predominantly located within the buried core of the protein. In these buried regions, pathogenic effects are approximately twice as likely to be caused by non-conservative amino acid substitutions as by conservative ones. This suggests that roughly 34–35% of pathogenic mutations in buried residues result from conservative substitutions, while the remaining 65–66% stem from non-conservative changes. A minor proportion of exposed residues were also observed in the pathogenic category with prevalence of non-conservative mutation. We found similar pattern of transition type distribution in the LD class. Conversely, the LT class shows entirely opposite distribution. Majority of LT is exposed and prevalent in non-conservative (55-62%) mutation. However, buried LT is predominant in conservative mutation (63-68%).

Missense mutation shows diverse conservative and non-conservative transitions of amino acids in benign and pathogenic variants. As the name suggests, non-conservative mutations change the characteristics of the residue. In line with the disruptive nature of non-conservative mutations, it predominates the pathogenic variant class, regardless of whether the residues are buried in the core or solvent-exposed (Figure S4). This pattern remains consistent for variants classified as LD, which shows a high prevalence across both structural states and solvent exposure (Figure S5). One of the limitations with our data is fewer number of ClinVar benign. That limits our result comparison with the benign. However, our results suggest that conservative mutation predominates in LT when buried and conversely, non-conservative mutations predominate the LT when exposed. Structural transitions move proteins through states stabilized by crucial interactions. Substantial changes in the stabilizing interactions (increase or decrease) can disrupt the dynamics of open and closed state. Variations in those residues often lead to pathogenic conditions. In our dataset, WT residues corresponding to 19 variations showed significant solvent exposure change. Among them 10 variants are non-conservative mutation (A737T, D743N, D743H, A747T, S813L, E1056K, K1227E, A1231T, G1232S, R1244C) and 9 are conservative mutation (D743E, A747V, F1052L, L1161F, N1208S, K1213R, R1244H, A1249V, P1289L). Thus, our results are consistent with the known physicochemical property shifts seen in pathogenic variants. Furthermore, the model identifies distinct amino acid transition patterns for likely tolerant variants and highlights residues essential for functional dynamics. This underscores the synergistic value of integrating structural scores with a mechanism-informed approach.

## Discussion

Precise and accurate prediction of variant effects remains a critical gap in the clinical research and implementation for patients with rare and undiagnosed diseases and cancers. Current genomics tools are constructed to maximize average performance across the genome, leading to their generic utility while also missing a level of domain-specific effects that are required for many disease-associated variants. Further, the naturally low frequency of individual rare disease cases restricts the data available for model training. In the case of SMARCA4, whose variation defines specific neurodevelopmental rare diseases and human cancers, only 10.3% of the unique missense changes are clinically dispositioned. This reality significantly limits the impact that genomic testing can have on patients, for providers, and towards therapeutic investigations, leading to the current unmet need for more accurate and mechanistic variant analysis approaches.

In this work, we demonstrate the potential for features derived from molecular mechanisms to augment conventional sequence-based features that effectively classify genetic variations. We applied unsupervised machine learning to implement a consensus model with 100% recall for pathogenic variants to be damaging and by leveraging a unique combination of evolutionary, structural, and mechanistic information. The most insightful observation of this work is to highlight residues that might play critical roles for transition between nucleotide-dependent states. Thus, such synthesis of features results in high-fidelity representation of the variant functional state which can be implemented for variant effect prediction with higher accuracy. Methods of the type we developed here have huge potential for helping to solve the challenge of interpreting human variation across rare and undiagnosed diseases where the case-specific data is limited but protein-specific data is primed for implementation.

The compact clustering of VUS we predict as LD around pathogenic and VUS we predict as LB around benign variation can be understood according to trends in their physical and chemical characteristics. The LT cluster is notably more compact than the LD cluster, suggesting a more restricted landscape of alternative amino acids that maintain a benign phenotype. This compactness indicates a prevalence of conservative substitutions that preserve the protein’s original physicochemical properties. In contrast, the LD cluster exhibits greater dispersion, likely driven by a higher proportion of non-conservative variations compared to LT variations, allowing for a more diverse and potentially destabilizing range of amino acid substitutions (Figure S5). This is further complemented by the result that there is a significant enrichment of buried residues, conserved residues and interfacial residues in LD, suggesting that residues that are conserved during evolution have critical role in the scaffold formation and preservation of interactions at the interface that might have transitory but crucial role for the state transformation, hence the mechanistic cycle. LB clusters on the other hand are enriched by surface exposed residues which suggest that if these residues are not involved in crucial interactions, then conservative mutations might not change their less critical roles. The nature of conservative versus non-conservative variation is also critical for the field. There is a strong tendency among geneticists to consider conservative variation as less likely to be pathogenic. In general, we observe this trend, but the exceptions are numerous. In fact, about one third of pathogenic and LD variation is conservative and about one third of tolerated and LB variation is non-conservative. Our findings underscore how more detailed modeling will produce more nuanced and context-specific classifications.

The strongest insights from this work is to identify residues that likely play critical roles in the transition between conformational states of the helicase. Our evaluation of solvent exposure and residue location identified nine residues with vital and differing properties for helicase transitions between ATP-bound and Apo states. Given their pathogenic status in ClinVar, these residues likely damage function, consistent with a fundamental role in the helicase mechanism. We find that mutations affecting the local stability of the domain’s core are typically non-conservative substitutions, yet about a third of them are conservative. Conversely, surface residues that are not involved in critical interactions are generally more tolerant of non-conservative variation. Thus, computational structural genomics methods are critical for the field to progress beyond site-specific scores and implement topology-dependent strategies.

Two primary limitations apply to this work. First, our integrated approach, which combines genomic pathogenicity scores with engineered structural and mechanistic calculations, is constrained by the small number of known benign variants. This bottleneck restricted our confidence in characterizing LB variation and led us to an all-versus-all analysis. Second, advanced variant effect assessment of this kind requires 3D protein models in multiple conformational states, and such structures are not always available for the protein of interest. We leveraged two for the current work, yet there are likely additional conformations of SMARCA4 that will be identified through future experiments and molecular dynamics simulations. Despite these constraints, the approach successfully characterized SMARCA4 helicase VUS, and the significant associations between our predictions and functional metrics provide independent validation of the results. Further, we assessed effects at the level of protein structure, leaving effects of DNA changes on gene expression, mRNA structure, etc., for future improvements. The methodology addresses a standing gap in genetic variant interpretation and offers a practical route to a clinical challenge specific to this disease-causing and druggable protein.

## Conclusion

Developing protein- or domain-specific variant effect prediction models faces annotation data scarcity and quality barriers. To overcome this challenge, we implemented a clustering-based approach to group variants with similar quantitative features and analyzed the consensus set that co-clustered with known benign and pathogenic variants. Our framework integrates structure and mechanism informed features with generic genome-wide sequence-based pathogenicity scores to develop the SMARCA4 helicase specific classification model. The method reclassified ClinVar VUS and CCP into LD and LB categories. LD clusters captured 100% of known pathogenic variants, and LB clusters captured 75% of known benign variants. The latter is notable: these benign variants sit outside the SNF2 and HelicaseC domains yet still share enough feature-level similarity to group with the LB set. Statistical enrichment and depletion of functional metrics across the reclassified variants provide independent support for these assignments. Thus, we have developed a protein-specific, mechanism-informed VEP for SMARCA4 helicase. The same framework extends to other proteins and enzymes where annotated data is limited, with direct relevance to undiagnosed disease and malignancy.

## Acknowledgments

Research reported in this publication was supported by the National Institute of General Medical Sciences of the National Institutes of Health under Award Number R35GM153740. The content is solely the responsibility of the authors and does not necessarily represent the official views of the National Institutes of Health. This research was completed in part with computational resources provided by the Research Computing Center at the Medical College of Wisconsin.

---

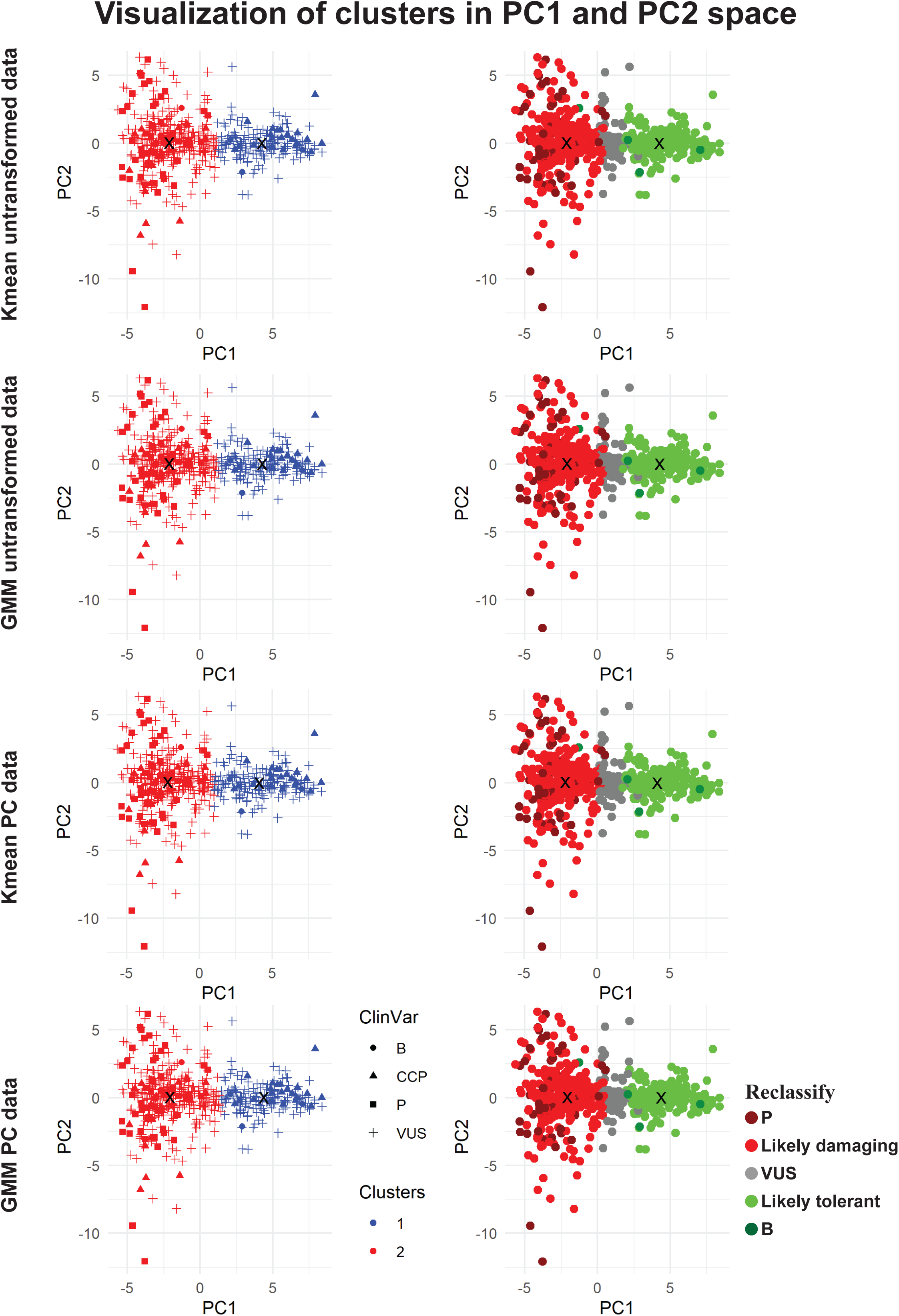

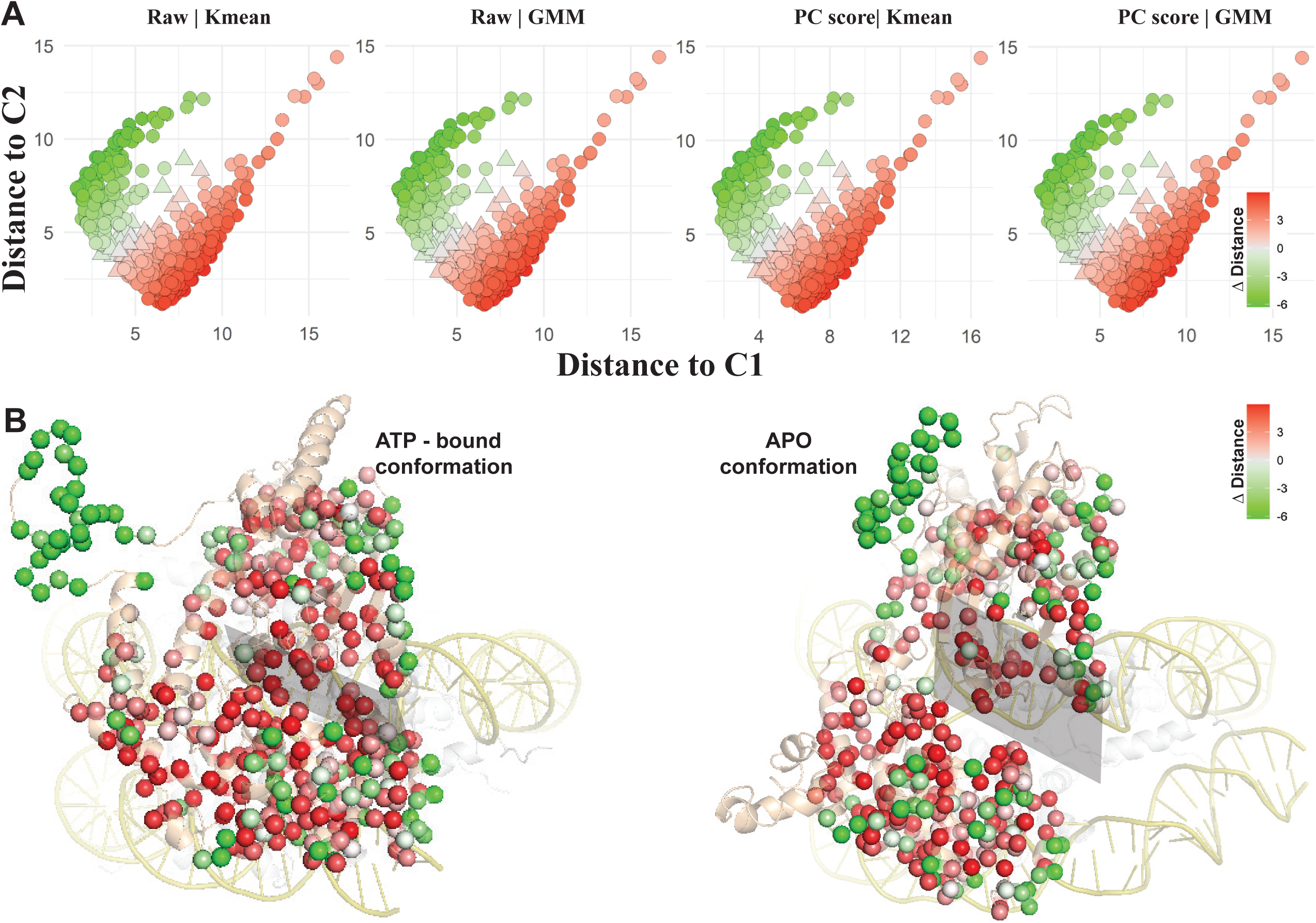

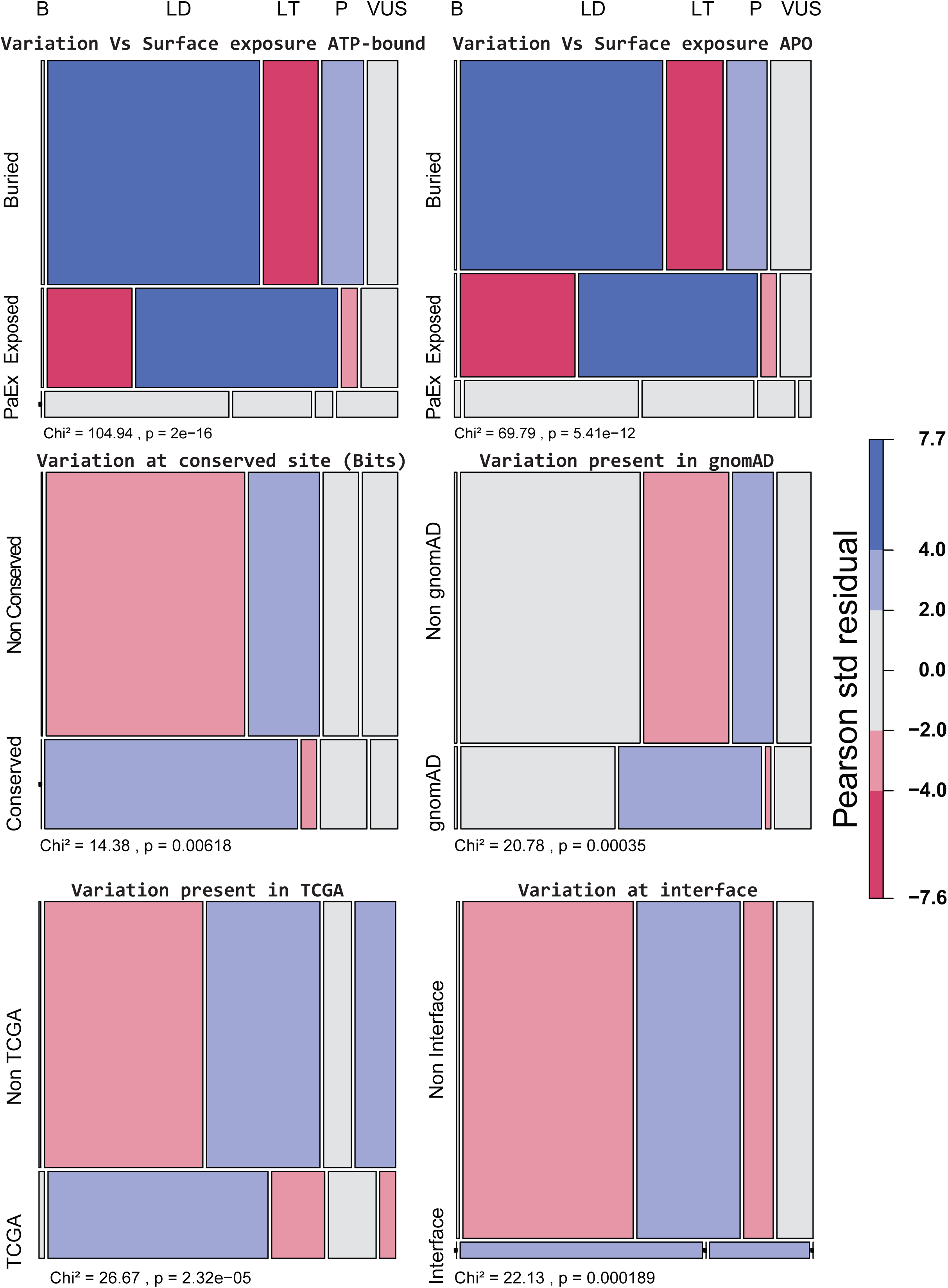

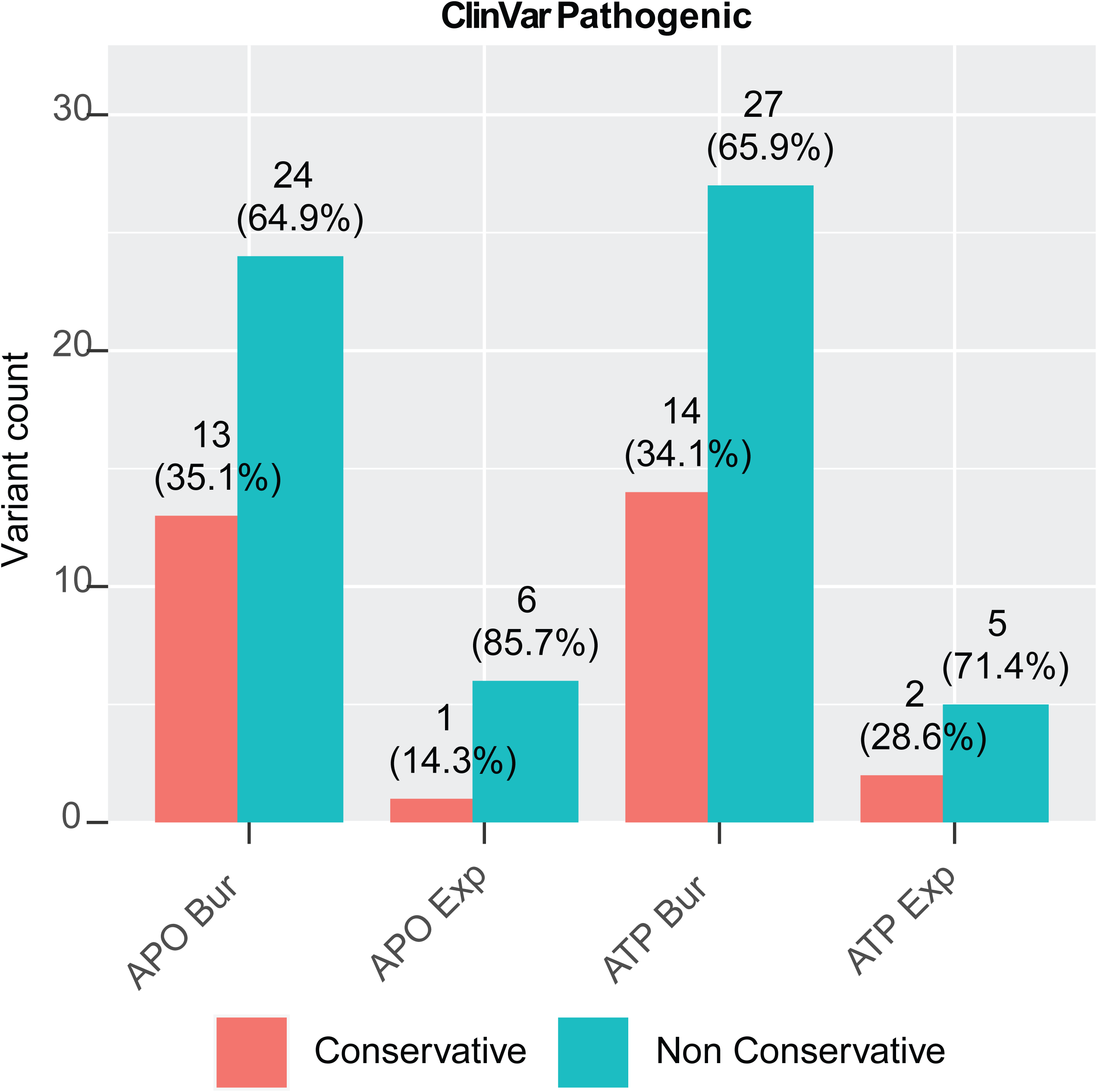

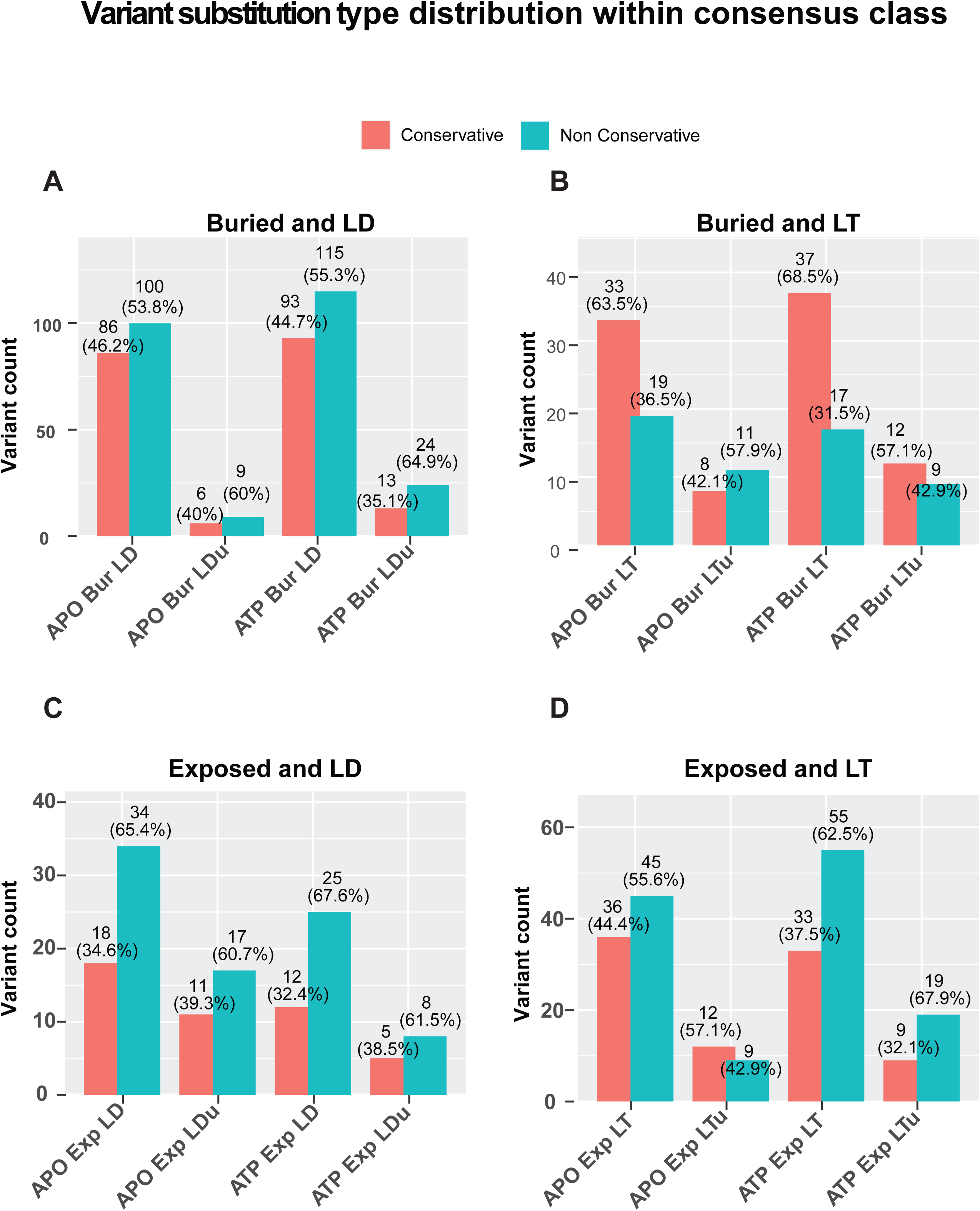

