## Supplementary figures and tables for "Defining the molecular tolerance-to-damage landscape of SMARCA4 helicase genetic alterations"

**Figure S1 (complementing Figure 3 and 4): ClinVar class, clusters, and reclassification class mapped on PC1 and PC2 for visualization.**

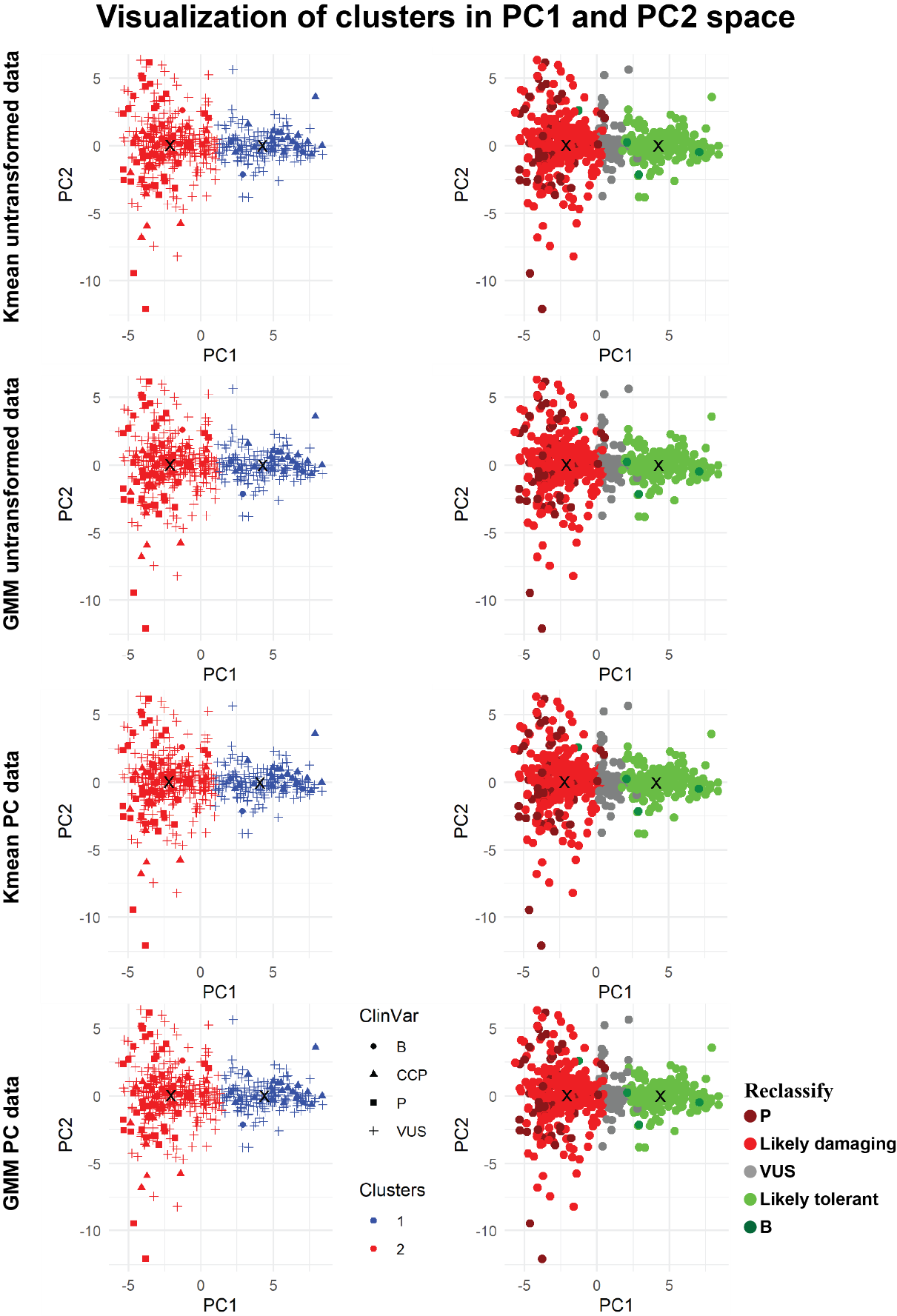

**Figure S2: A)** Distance difference plot from each centroid. Variants remaining as VUS are shown as gray triangles, green circles are Likely tolerant, and red circles are Likely damaging. **B)** The same data are projected onto the ATP-bound conformation (PDB: 7VDV) and **C)** the unliganded (a.k.a., Apo) conformation (PDB: 9A0K).

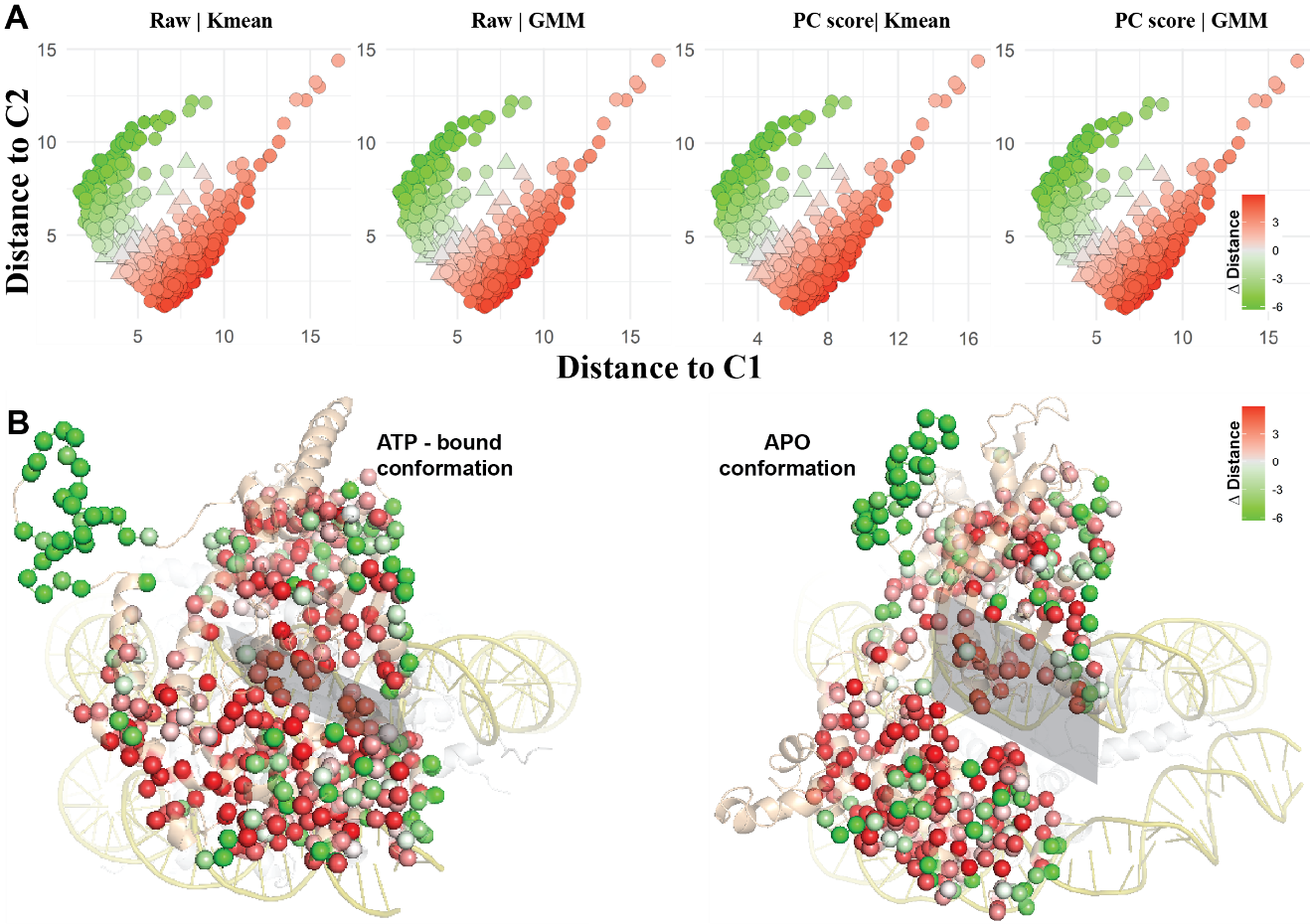

**Figure S3: Enrichment of functional metrics validates our consensus classification**: The panel of six figures shows significant enrichment of functional metrices with known and predicted class, such as, buried (B), likely damaging (LD), likely tolerant (LT), pathogenic (P) and VUS, shown on the top of the first row.

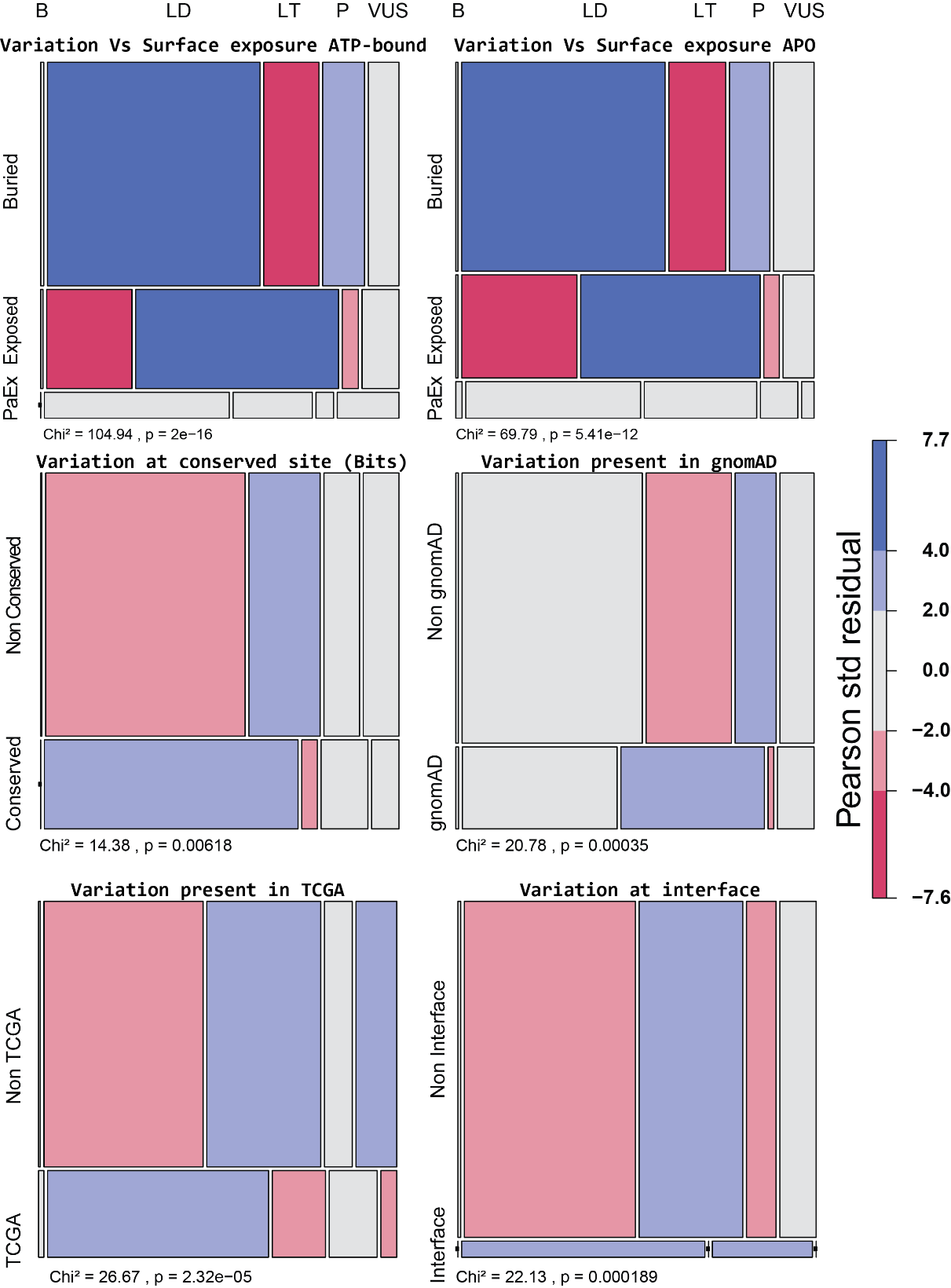

**Figure S4: Change of physicochemical properties on residue substitution** show distinct distribution in buried (Bur) and exposed (Exp) residues of missense mutation.

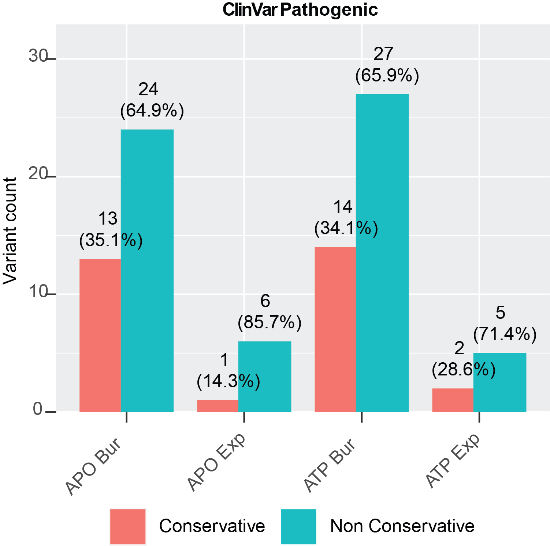

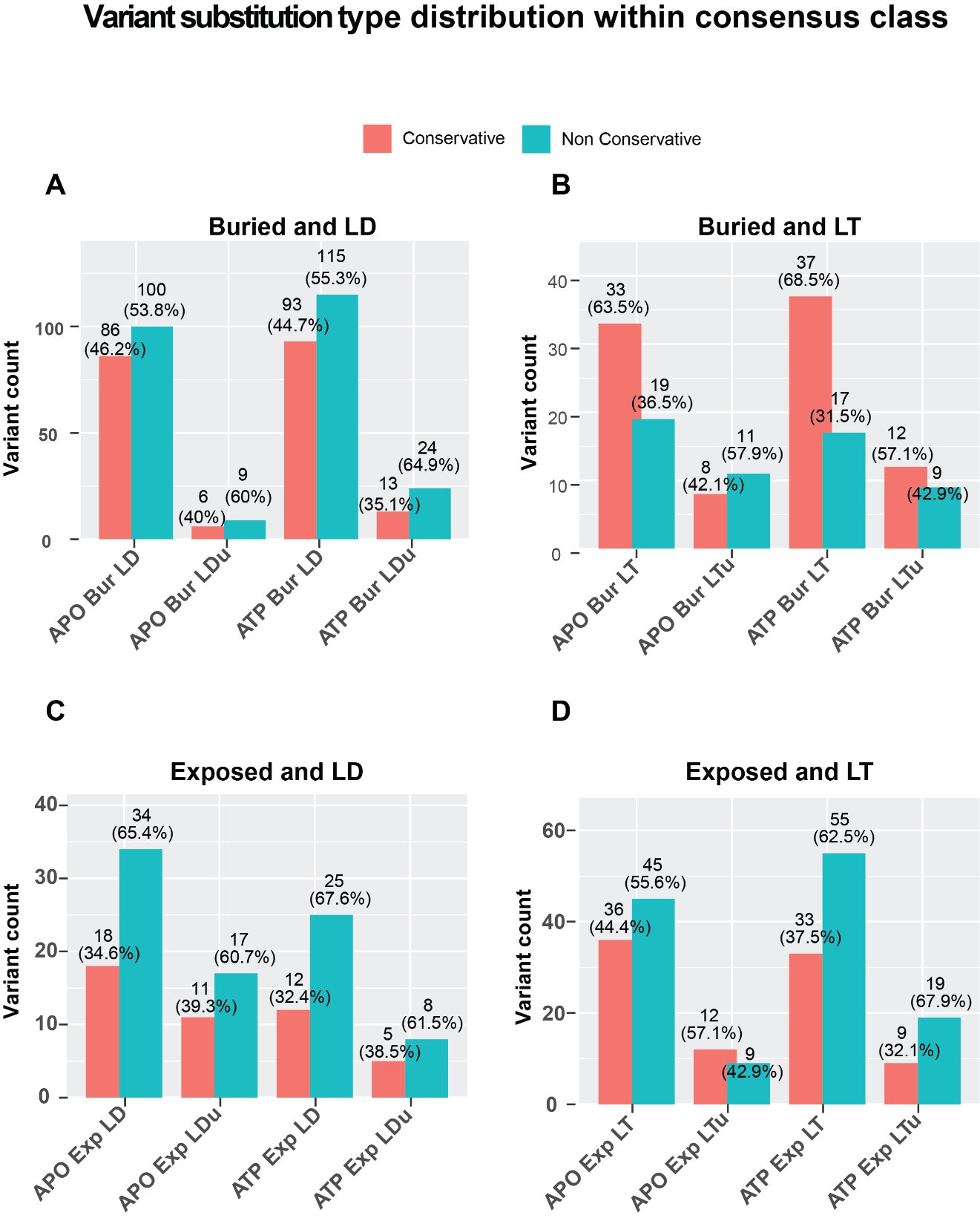

**Figure S5: Consensus class distribution of mutation types (conservative and non-conservative):** Distribution of missense mutation type in consensus classes follows similar pattern as observed in pathogenic mutation. ATP and Apo states show similar distribution in Exposed-LD/LT and buried-LD except buried-LT. Mutation unique to certain category is show by ‘u’ at the end.

**Table S1: Significant enrichment of functional metrics validates the results:** Statistical test results, such as Pearson Chi-square, Likelihood ratio and Fishers’s exact test, for all functional metrics are shown with their p-values.

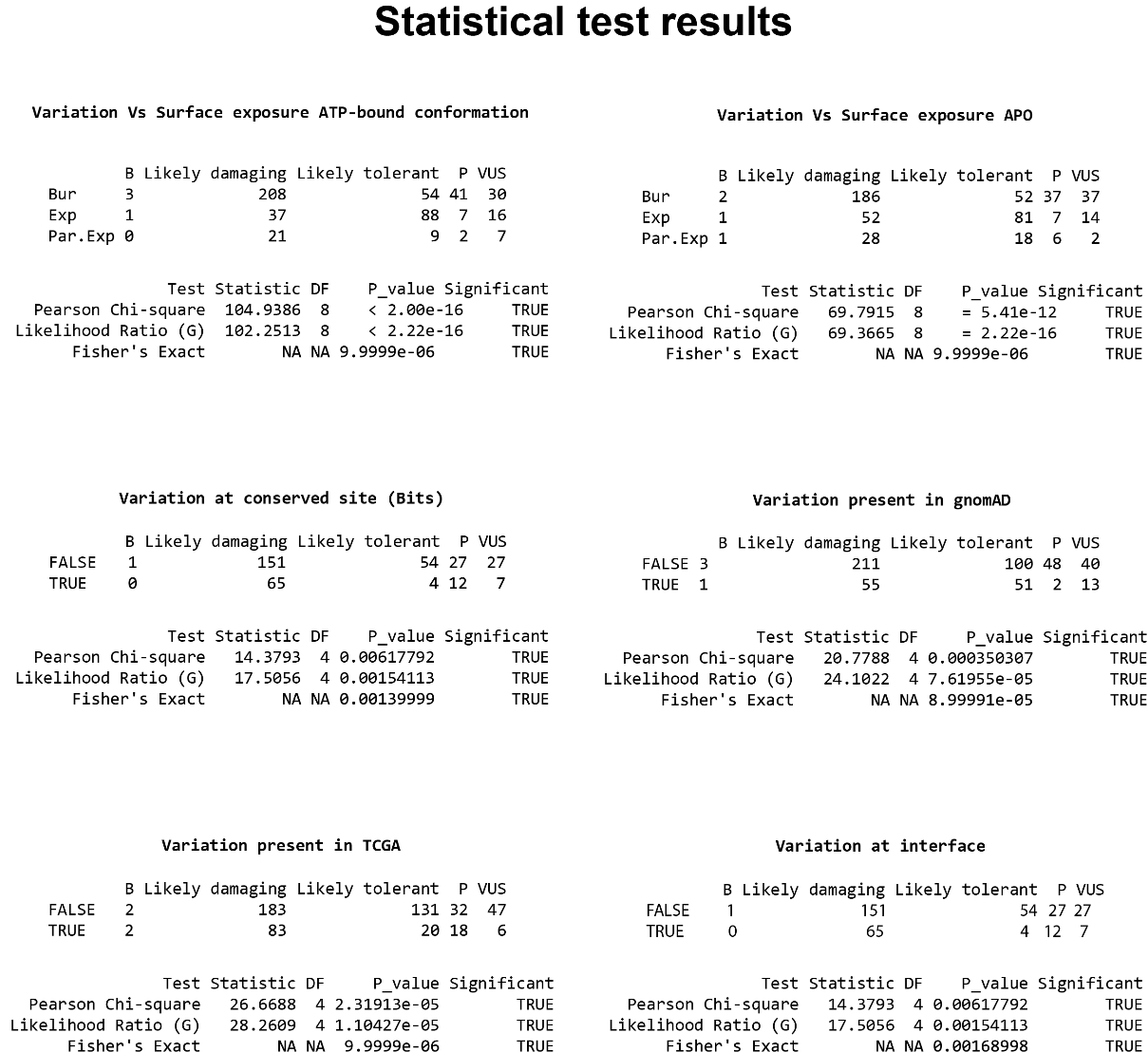
